# Description, taxonomy, and comparative genomics of a novel *Thermoleptolyngbya* strain isolated from hot springs of Ganzi, Sichuan China

**DOI:** 10.1101/2021.01.08.426015

**Authors:** Jie Tang, Liheng Li, Meijin Li, Lianming Du, Michał Waleron, Małgorzata Waleron, Krzysztof Waleron, Maurycy Daroch

## Abstract

*Thermoleptolyngbya* is a newly proposed genus of thermophilic cyanobacteria that are often abundant in thermal environments. However, a vast majority of *Thermoleptolyngbya* strains were not systematically identified, and genomic features of this genus are also sparse. Here, polyphasic approaches were employed to identify a thermophilic *Thermoleptolyngbya* A183 (FACHB-2491) isolated from hot spring Erdaoqiao, Ganzi prefecture, China. Morphological characterization was investigated by microscopy. Whilst the results of 16S rRNA were inconclusive, the phylogenetic reconstruction and secondary structures of 16S-23S ITS strongly suggested that A183 strain is a novel species within *Thermoleptolyngbya*. Moreover, genome-scale average nucleotide identity (ANI) confirmed the genetic divergence of A183 from *Thermoleptolyngbya* O-77. Comparative genome analysis revealed inconsistent genome structures of *Thermoleptolyngbya* strains. Further GO analysis showed that the specific genes were distributed in a wide range of functional categories. In addition, analysis of genes related to thermotolerance, signal transduction, and carbon/nitrogen/sulfur assimilation revealed the ability of this strain to adapt to inhospitable niches in hot springs.

## 1 Introduction

Cyanobacteria are widely distributed microorganisms in various ecological niches due to abundant features allowing extensive adaptations. In particular, cyanobacterial populations existing in thermal environments have attracted increased interests in light of a crucial role in energy metabolism and matter cycling in ecosystems (Amin et al., 2017). Among the cyanobacteria communities, strains morphologically assigned to genus *Leptolyngbya* are often reported to be prosperous in many thermal environments (Mackenzie et al., 2012; Amarouche-Yala et al., 2014; Tang et al., 2018a; Strunecký et al., 2019).

Identification of *Leptolyngbya*-like strains has been controversial because of their simple morphology, lacking significant discrimination. The heterogeneity of *Leptolyngbya* has been questioned since the establishment of this genus (Bruno et al., 2009). The genus *Leptolyngbya* has been recognized as polyphyletic (Johansen et al., 2011; Perkerson et al., 2011), and there are strong recommendations to conduct a taxonomic reevaluation of this genus. In light of limited information provided by cell morphology investigations of trichomes, genetic and molecular techniques have been applied to facilitate the establishment of correct taxonomy. The 16S rRNA gene has been proposed as a universal DNA barcoding marker for species-level identification of bacterial isolates, and as a complement to morphology-based taxa identification. However, in some cases, the 16S rRNA gene cannot resolve cyanobacterial phylogeny at the species level (Niclas et al., 2010; Eckert et al., 2014). Additional genomic locus, 16S-23S intergenic spacer (ITS), has been used for cyanobacterial systematics. It has been applied for the construction of phylogenetic trees and through the analysis of secondary structures of 16S-23S ITS regions (Johansen et al., 2011). Numerous studies confirmed the integrated approach of analyzing 16S rRNA gene phylogeny and 16S-23S ITS secondary structure to be useful and robust for complex cyanobacterial taxonomy, as in the case of the species or genera within the family Leptolyngbyaceae (Komarek et al., 2014; Debnath et al., 2017; Shalygin et al., 2020). Recently, *Thermoleptolyngbya*, a cryptogenus newly delineated using 16S rRNA gene and 16S-23S ITS, emerged from strains originally grouped into *Leptolyngbya* (Sciuto & Moro, 2016). Two species were ascribed to this new taxon: *T. albertanoae* and *T. oregonensis*. Still, the majority of strains ascribed to *Thermoleptolyngbya* were not systematically identified, and 16S rRNA gene sequences were exclusively used for their taxonomic recognition. As such, the genus requires expanded sequence information to complement the 16S rRNA gene taxonomy by multiple-locus sequence analysis (MLSA) or on the whole genome level. The widened sequence space, especially about the underrepresented members of the genus, is required to guide phylogenetic and taxonomic studies and eventual reclassification. In addition, acquisition of complete genome may provide new insights into the genomic features of genus *Thermoleptolyngbya*, particularly the survival mechanism in thermal conditions from a genomic perspective.

In the current study, a *Thermoleptolyngbya*-like strain, A183 originally isolated from Erdaoqiao hot springs (Tang et al., 2018a) in Ganzi Prefecture, Sichuan Province, China, and as described previously (Tang et al., 2018b), was used for taxonomic reevaluation. The morphological and molecular data were collected for this thermophilic *Thermoleptolyngbya*. This work aimed to provide deeper insights into taxonomy and genomic features of *Thermoleptolyngbya* strains. Characterization was achieved by analysis of its morphology, 16S rRNA/ITS sequence, secondary structures of ITS regions and the genome.

## 2 Material and Methods

### 2.1 Information on cyanobacterial strain A183

The stock culture of strain A183, cryopreserved for over two years as 10% DMSO in BG11 stock in −80°C, was used to establish the final precultures for experiments essentially as described by Tang et al. (2018b). The shake-flask cultures were grown in a shaking incubator in BG11 medium at 45 °C under a photoperiod of 16-h light (45 μmol m^-2^ s^-1^) and 8-h dark. The strain initially denoted, and deposited in Peking University Algae Collection as PKUAC-SCTA183 has been also deposited in the Freshwater Algae Culture Collection at the Institute of Hydrobiology (FACHB-collection) with accession number FACHB-2491.

### 2.2 Genome sequencing and assembly

Genomic DNA of strain A183 was extracted and purified using a bacterial genomic DNA isolation kit (Generay, Shanghai, China) according to the manufacturer’s instructions. Purified genomic DNA was subjected to gel electrophoresis and spectrophotometric measurements for quality and quantity assessment, respectively. The whole-genome sequencing of A183 was performed using a hybrid sequencing strategy combining PacBio long reads and Illumina short reads. Two SMRT cells were used for PacBio sequencing and yielded 57,735 adapter-trimmed reads (subreads) with an average read length of approximately 9 kb, corresponding to 94-fold coverage. Illumina NovaSeq sequencing of A183 generated a total of 6,810,074 filtered paired-end reads (clean data), providing approximately 185-fold coverage of the genome. These clean data were assembled into contigs using MaSuRCA v. 3.3.9 with default parameters (Zimin et al., 2013), finally generating a single contig. The genome obtained was mapped with Illumina reads using BWA v0.7.17 (Li & Durbin, 2009) and then Pilon v1.23 (Walker et al., 2014) to correct any assembly and sequence errors. The complete genome has been deposited in GenBank with an accession number CP053661.

### 2.3 Genome annotation and comparative genomes analysis

The genome of A183 was annotated using a modified pipeline previously established by Tang et al. (2019). Briefly, gene prediction and annotation were automatically performed using the NCBI prokaryotic genome annotation pipeline (Pruitt et al., 2009), and further using the RAST annotation system to minimize poor calls. The insertion sequence (IS) was detected and annotated by ISsaga (Varani et al., 2011). Prophage regions were predicted by PHASTER (Arndt et al., 2016). CRISPR loci were detected using CRISPRCasFinder server (Grissa et al., 2007). The protein sequences predicted by RAST were aligned against the NCBI non-redundant database using BLASTP with an *E*-value cut-off of 1e-5. The alignment results were imported into BLAST2GO V5.2.5 (Conesa et al., 2005) for GO term mapping. The results of BLAST2GO analysis were submitted to the WEGO (Ye et al., 2006) for GO classification under the biological process, molecular function and cellular component ontologies. The circular plot of A183 genome was generated by Circos v0.68 (Krzywinski et al., 2009).

The genome of *Thermoleptolyngbya* strain O-77 (AP017367) was used for comparative genomic analysis with strain A183 (CP053661). The genome sequence was also subjected to the annotation pipeline mentioned above to keep all the data analyzed under the same criteria. To compare the gene context, all-against-all BLASTP alignments were performed using the following thresholds, E-value cut-off of 1E-5 and 30% identity and the best hit of alignments were selected. The average nucleotide identity (ANI) and the percentage of genome segments mapped were calculated using FastANI (Jain & Rodriguez, 2018).

### 2.4 Phylogenetic reconstruction

Sequences of the 16S rRNA gene and 16S-23S ITS were extracted from A183 genome for phylogenetic analysis. Reference sequences of cyanobacteria were also retrieved from GenBank through BLAST search for 16S rRNA gene and 16S-23S ITS dataset construction, respectively. Multiple alignments of sequences were generated with Muscle incorporated in Mega7 (Kumar et al., 2016). Alignments were subjected to manual editing where necessary. Sequences of each alignment were trimmed to the same length.

Phylogenetic trees of 16S rRNA and 16S-23S ITS sequence datasets were reconstructed using Maximum-Likelihood (ML), Maximum-Parsimony (MP), and Neighbor-Joining (NJ) methods, respectively. ML phylogenetic analyses were both carried out using PhyML version 3.0 (Guindon et al., 2010) and the substitution models were selected based on the Akaike information criterion (AIC) by Model Selection function implemented in PhyML (Vincent et al., 2017). The NJ trees were both constructed using General Time Reversible (GTR) model implemented in Mega7. Nonparametric bootstrap tests (1000 replications) were applied to evaluate the robustness of tree topologies.

### 2.5 Secondary structure Prediction

The tRNAs presented in 16S-23S ITS sequences were predicted by tRNAscan-SE v1.3.1 (Lowe & Eddy, 1997). The conserved domains (D1-D1’, D2, D3, boxA, and D4) and the variable regions (V2, boxB, and V3) of 16S-23S ITS were detected as reported by Iteman et al. (2000). The secondary structures of these DNA fragments were individually folded by Mfold web server (Zuker, 2003). Except for the use of the structure draw mode untangle with loop fix, default conditions in Mfold were used in all cases.

### 2.6 Microscopic analysis

The isolated cyanobacterium was investigated using light microscopy (LM, DP72; OLYMPUS, Tokyo, Japan). Approximately 50 ul of culture was dropped on the microscopy slide and observed under 400x magnification. The images were captured using U-TV0.63XC camera (OLYMPUS, Tokyo, Japan). Scanning electron microscopy (SEM) was performed as follows. Cells were washed gently with PBS (Servicebio, G0002), and fixed for 2 hours in fixation solution (Servicebio, G1102). Subsequently the cells were post fixed with 1% OsO_4_ (TedPellaInc) in 0.1 M PB (pH 7.4) for 1-2 h at room temperature. The fixed material was dehydrated in a graded ethanol series (30% to 100%) (Sinopharm) and isoamyl acetate (Sinopharm) for 15 min and dried with Critical Point Dryer (Quorum, K850). Then specimens were attached to the metallic stubs using carbon stickers and sputter-coated with gold for 30 s. Coated samples were examined directly under the scanning electron microscope (HITACHI, SU8100). For transmission electron microscopy (TEM), the fixation solution (Servicebio, G1102) was added to the isolated cells. The cells were subsequently pelleted and resuspended in the fresh fixation solution. Cooled 1% agarose solution was mixed with the cells and the agarose blocks were post fixed with 1% OsO_4_ in 0.1 M PB (pH 7.4) for 2 h. Then cells were dehydrated as described above for SEM, and embedded in pure EMBed 812 resins (SPI, 90529-77-4). Embedded cells were incubated in 65°C oven for more than 48 h to complete polymerization. The sections were cut to 60-80 nm thin layers using the ultra-microtome (Leica, LeicaUC7), stained with 2% uranium acetate saturated alcohol solution and lead citrate for 8 min, and examined using TEM (HITACHI, HT7800).

## 3 Results and discussion

### 3.1 General features of strain A183 (FACHB-2491) genome

The combined assembly of PacBio and Illumina sequencing data successfully generated the complete genome of strain A183. The genome (Fig. 1) comprises a single circular chromosome with a size of 5,525,100 bp (GC content, 56.38%) and no plasmid. Gene prediction and annotation of strain A183 resulted in 5,166 protein-coding sequences (CDS), approximate half (50.3%) of which were identified as hypothetical proteins. Functional distribution on GO categories of all CDS identified was summarized in Fig. S1. Two ribosomal RNA (*rrn*) operons were detected and 45 tRNA genes were predicted in the A183 chromosome (Table 1).

**Fig. 1.**
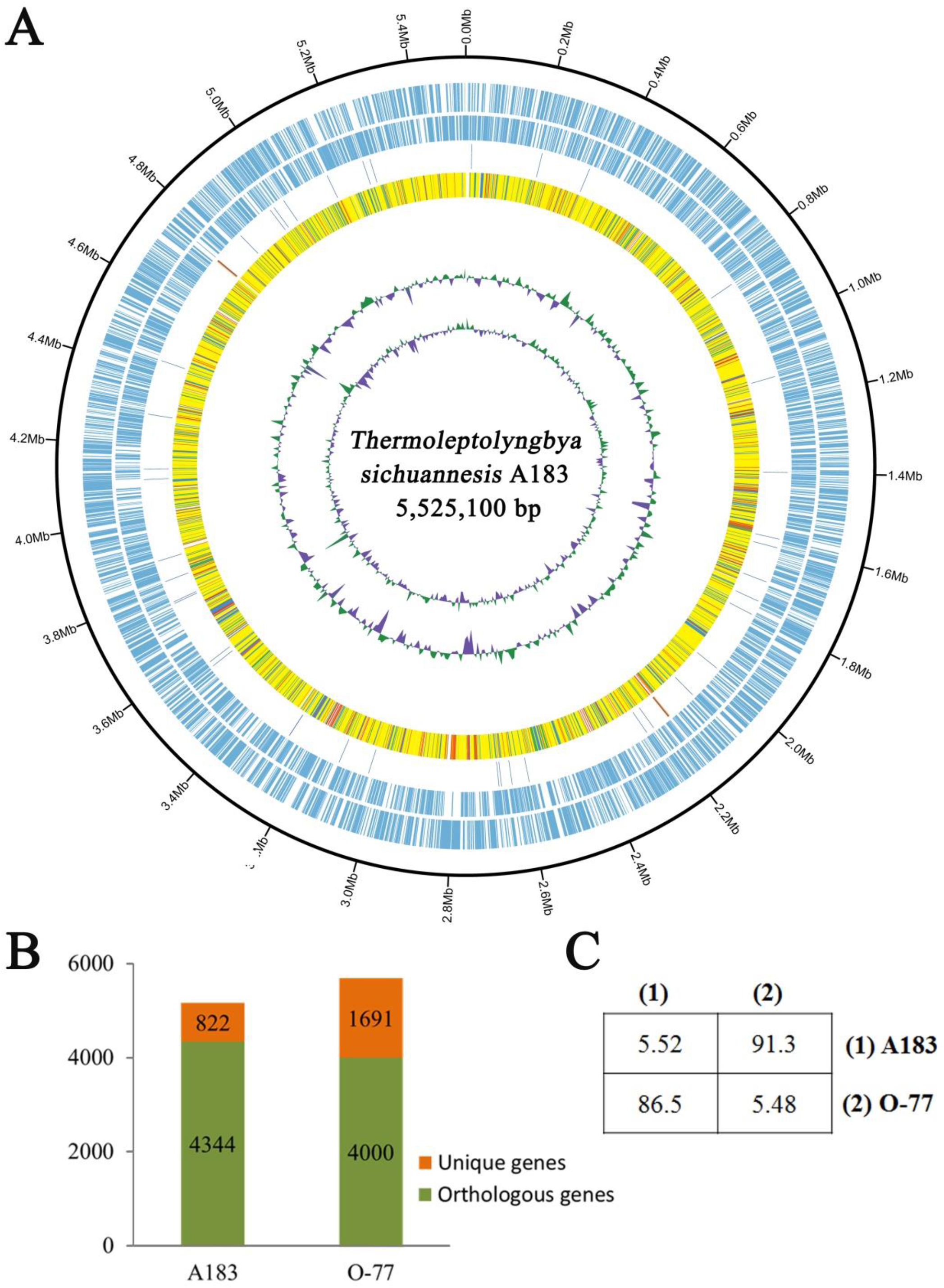
Comparison between *Thermoleptolyngbya* strains. **A**: circular plot of A183 genome. Rings are as follows (outer - inner): CDS on plus strand; CDS on minus strand; rRNA (orange) and tRNA (blue); the shared amino acid identities of BLASTP alignments with O-77; the last two circles represent GC content and GC skew both calculated for a 10-kb window with 1-kb stepping. The colour scheme for the heat map of orthologs is as follows: yellow, orthologs ≥ 90% identity; blue, 80 - 90% identity; green, 70 - 80% identity; red, 50 - 70% identity; orange, 30 - 50% identity. **B**: number of shared and specific genes between strains. **C**: Pairwise genome sequence similarity scores. The numbers along the diagonal, above diagonal and below the diagonal referred to the genome sizes (Mb), the average nucleotide identity (ANI) values (%) and the proportion of segments that could be mapped between the two genomes for ANI calculation, respectively.

**Table 1.**
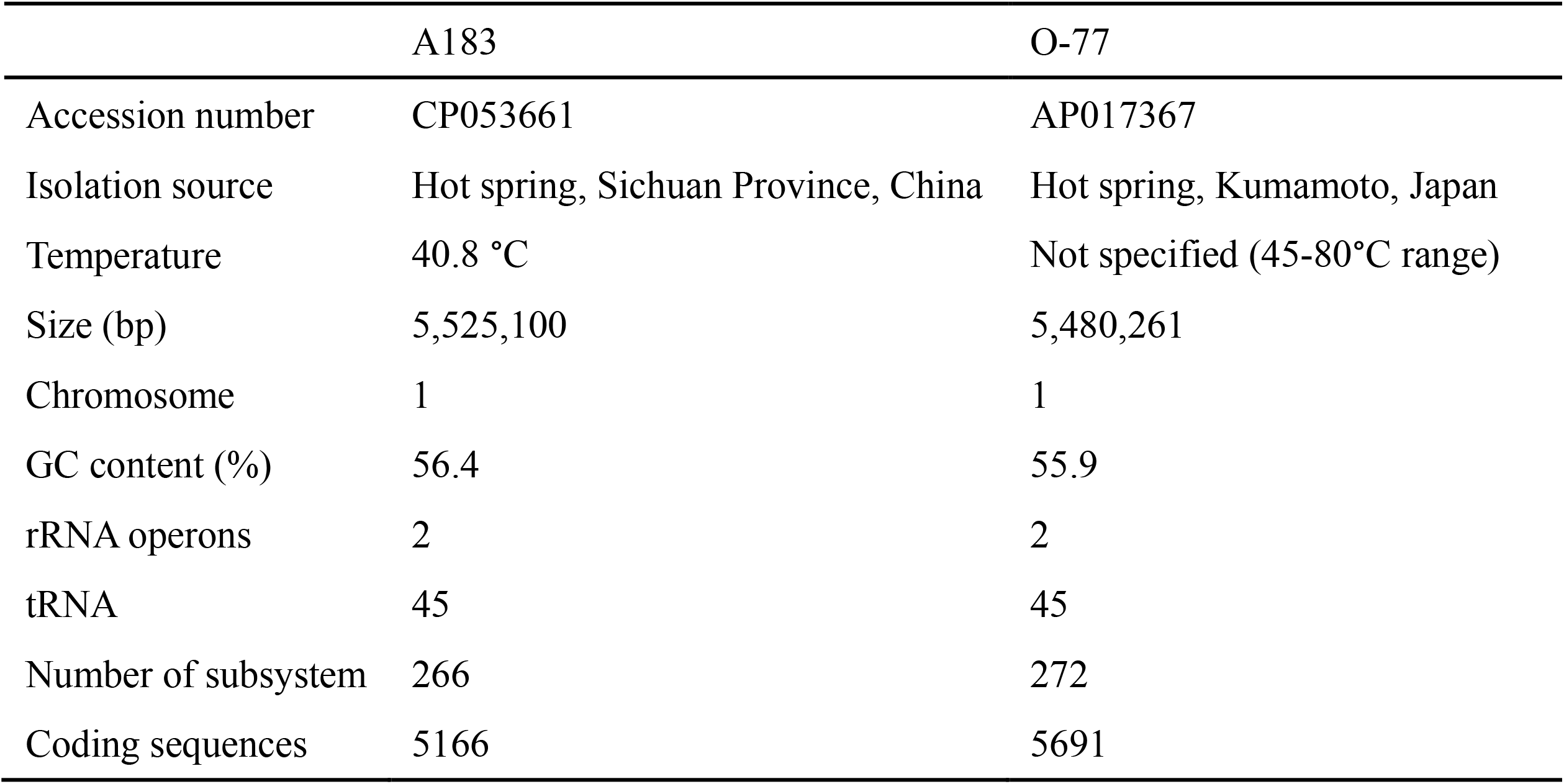
Genome features of *Thermoleptolyngbya* strains A183 and O-77. NA: not available

In the A183 chromosome, 186 ISs representing 28 different ISs were identified. The most frequently observed IS type was the ISKra4 family (30.11%), followed by IS630 family (26.88%) and IS4 family (19.89%). Numerous genes encoding transposase (Table S1) were also observed, indicating that the genetic plasticity of the strain might be shaped by intragenomic rearrangements. It was proposed that transpositions play a crucial role in genomic rearrangements and are involved in gene regulation and adaptation processes that determine the directions of microevolutionary processes in cyanobacteria (Mikheeva et al., 2013).

The A183 chromosome harboured around 190 transporter-related genes (Table S1). Among these transporters, ABC transporters accounted for the majority, and distinct bias was found in many genes, such as P-type ATPase transporter for copper which only has two copies. Functionally, these transporters have been predicted as Na^+^/H^+^, iron, phosphate, amino acid, bicarbonate, and CO_2_transporters etc.

### 3.2 Phylogeny of 16S rRNA

To ascertain the taxonomic position of strain A183, a ML phylogenetic tree was reconstructed based on 16S rRNA gene sequences of the 58 cyanobacterial strains. The ML tree (Fig. 2) resolved 13 well-defined clades of isolates corresponding to previously described genera: *Alkalinema* (Vaz et al., 2015), *Halomicronema* (Abed et al., 2002), *Haloleptolyngbya* (Pawan et al., 2012), *Kovacikia* (Miscoe et al., 2016), *Leptolyngbya sensu stricto* (Taton et al., 2010), *Nodosilinea* (Perkerson et al., 2011), *Oculatella* (Zammit et al., 2012), *Pantanalinema* (Vaz et al., 2015), *Phormidesmis* (Komárek et al., 2009), *Plectolyngbya* (Taton et al., 2011), *Stenomitos* (Miscoe et al., 2016), *Thermoleptolyngbya* (Sciuto & Moro, 2016), and *Gloeobacter* as an outgroup of the tree. Strain A183 investigated in this study closely clustered with 15 strains affiliated to genus *Thermoleptolyngbya*. Three clusters were not assigned to the previously described taxa and were marked as cluster A to C, respectively. In addition, the sequences of “*L. antarctica*” ANT.L67.1 and *L. indica* LKB did not collocate with any cluster and were placed in separate branches.

**Fig. 2.**
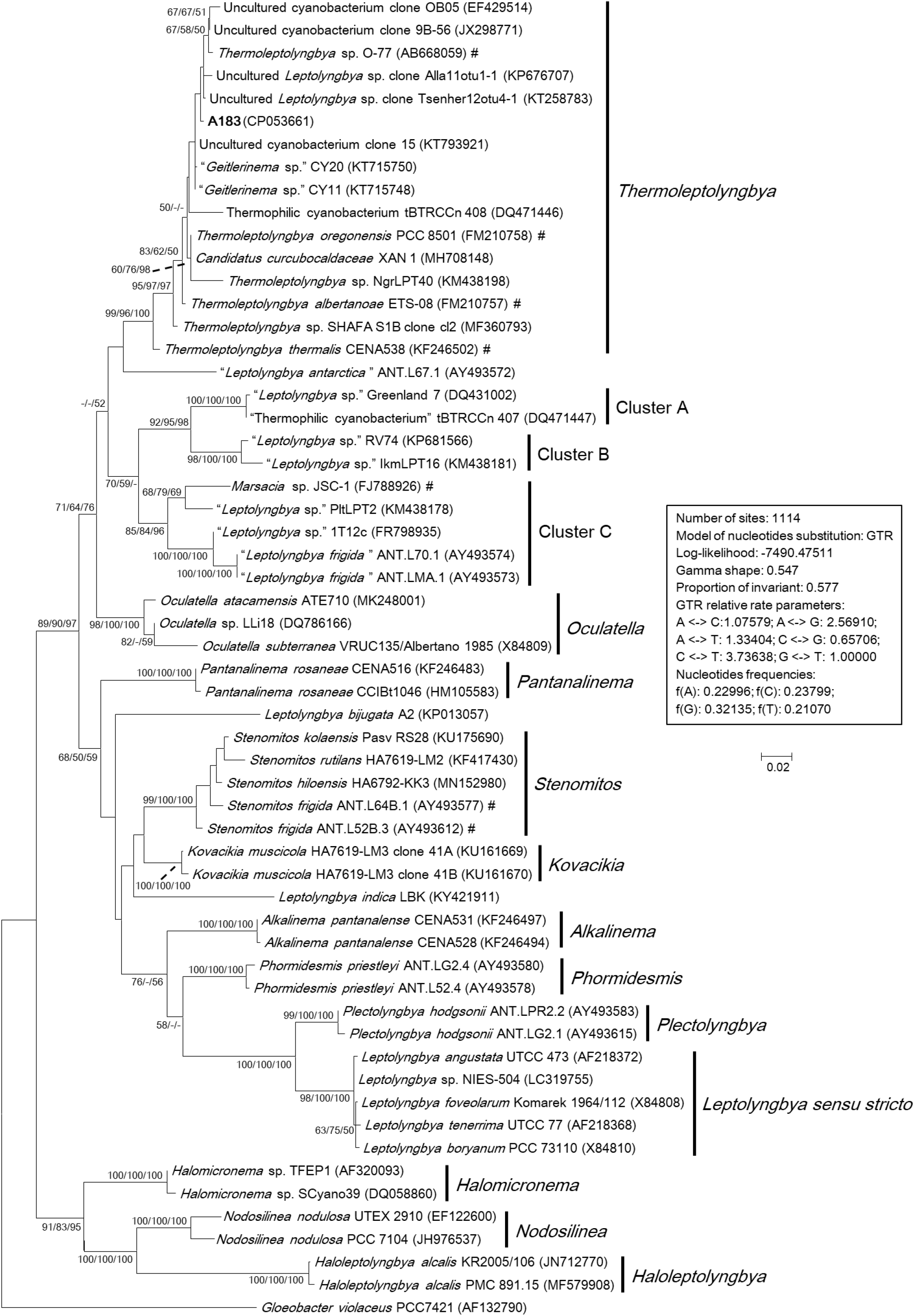
Maximum-likelihood phylogenetic tree of 16S rRNA gene sequences. The numbers at nodes refers to the support values of ML, MP, and NJ analysis, respectively. Only bootstrap values>50% (1000 nonparametric replications) are indicated at nodes. Genera or cluster are indicated by vertical bars. Scale bar = 2% substitutions per site. Strains in “quotation marks” have uncertain genus name. Strains marked with # have been recently reclassified or proposed to be reclassified.

The sequence identities of 16S rRNA gene were calculated between A183 and other strains phylogenetically grouped into *Thermoleptolyngbya* clade (Table S2). The A183 showed sequence identities ranging from 97.11 to 99.46%, compared to the other strains assigned to *Thermoleptolyngbya* clade. Notably, numerous strains within *Thermoleptolyngbya* clade were labeled by uncertain genus names (Fig. 2), while the sequence identities of 16S rRNA gene (Table S2) strongly indicated that these strains are members of genus *Thermoleptolyngbya*, according to the recommended threshold for bacterial species (98-99%) or genera (94.5-95%) demarcation (Rodriguez-R et al., 2018). Moreover, these *Thermoleptolyngbya* strains originated from thermal environments worldwide (Table S2) (Lacap et al., 2007; Oren et al., 2008; Peng et al., 2013; Nakamori et al., 2014; Gaisin et al., 2015; Bravakos et al., 2016; Sciuto & Moro, 2016; Heidari et al., 2018; Tang et al., 2018a), except for XAN 1 and CENA538. This is consistent with the general knowledge that organisms belonging to *Thermoleptolyngbya* appear to have thermal origins (Sciuto & Moro, 2016). Information on XAN 1 is sparse, and perhaps it was an inhabitant of hydrothermal spring based on the submission title on NCBI. CENA538 isolated from saline-alkaline Lake during the Brazilian dry season was subjected to dessication periods, and the temperature and high salinity of sampling site could be considered as a thermal environment (Andreote et al., 2014).

The results of phylogenetic reconstruction (Fig.2) and sequence identity (Table S2) indicated that 16S rRNA gene might not effectively differentiate species-level relationship of strains belonging to *Thermoleptolyngbya*. A verified example was the establishment of two different *Thermoleptolyngbya* species (*T. oregonensis* and *T. albertanoae*) (Sciuto & Moro, 2016), whereas the phylogenetic relationship (Fig. 2) and sequence identity (99.17%) of the two strains were susceptible to reach an erroneous conclusion on species differentiation. Therefore, molecular and phylogenetic analysis of more genomic loci is inordinately crucial for accurate species identification within *Thermoleptolyngbya*.

As expected, the strains affiliated to genus *Leptolyngbya* were widely distributed in the ML tree, suggesting interspecific heterogeneity within this genus. *L. indica* LBK and *L. bijugata* A2 have been recently proposed as new species of genus *Leptolyngbya* based on morphology and molecular data (Wang et al., 2015; Debnath et al., 2017). These findings are in according with their positions in the ML tree generated in this study. An emendation has been recently proposed on Antarctic strains previously reported as *“Leptolyngbya. frigida”* to *Stenomitos. frigidus* (Fig. 2) (Shalygin et al., 2020). These strains are likely to belong to newly reported genus *Stenomitos* isolated from cave habitats on Kauai, Hawaii (Miscoe et al., 2016). Another Antarctic strain ANT.L67.1 was also previously identified as “*L. antarctica*” (Taton et al., 2010), whereas the phylogenetic position (Fig. 2) and sequence identity (Table S2) indicated that ANT.L67.1 might putatively represent a new genus within family Oculatellaceae. Similarly, these so-called *Leptolyngbya* strains in Cluster A to C are phylogenetically suspicious to be new species or new genus within family Oculatellaceae.

Cluster A included two strains both from hot springs, namely “*Leptolyngbya* sp.” Greenland 7 from a microbial mat of Kap Tobin (Arctic hot spring, Greenland) and tBTRCCn407 from Zerka Ma’in spring in Jordan. Cluster B was formed by one unknown strain, “*Leptolyngbya* sp.” RV74, and “*Leptolyngbya* sp.” IkmLPT16, isolated from the Moustafa hot spring in Greece. The phylogenetic relationship of cluster A and B (Fig. 2) as well as 16S rRNA identity analysis (Table S2) suggested that the two clusters putatively represent two new genera.

Cluster C comprised five strains, two of which (PltLPT2 and JSC-1) were assigned to *Marsacia*, two Antarctic strains identified as “*L. frigida*”, and the last one named as “*Leptolyngbya* sp.” 1T12c. JSC-1 was polyphasically characterized as *M. ferruginose* (Brown et al., 2010), while information on *“Leptolyngbya* sp.*”* PltLPT2 is scarce and should be a sister taxon to JSC-1 based on the phylogenetic position (Fig. 2) and 16S rRNA identity (Table S2). The sequence identities (Table S2) among strains of Cluster C suggested that they belong to the same genus but different species, although proper genus naming has not been verified yet. Besides, the strong evolutionary relationship and 16S rRNA identity indicated that ANT.LMA.1, ANT.L70.1 and 1T12c putatively represent two species of the new genus.

The strong evolutionary relationship of the aforementioned *Leptolyngbya*-like strains (Fig. 2) showed apparent separation from *Leptolyngbya sensu stricto* and was also observed in previous studies (Isabella Moro & Andreoli, 2010; Roy et al., 2013; Sciuto & Moro, 2016). These potentially new phylotypes will need to be investigated carefully in future work using polyphasic techniques, such as ecological, morphological, and molecular data (MLSA or genome).

### 3.3 Phylogeny of 16S-23S ITS

It is well known that cyanobacterial strains could be divided into ambiguous or cryptic species or genera through molecular identification using 16S rRNA gene alone. Here, a multilocus approach was employed, and additional phylogenetic analysis was performed using 16S-23S ITS sequences since the ITS region have been commonly used to establish cyanobacterial ecotypes or species (Sciuto & Moro, 2016; Brito et al., 2017).

The phylogenetic reconstruction based on full-length 16S-23S ITS sequences (Fig. 3) was consistent with the phylogeny of the 16S rRNA gene, although fewer sequences were included in the analysis. The *Alkalinema* clade was rooted as an outgroup. Strains ascribable to *Thermoleptolyngbya* were placed into a well-supported clade and showed evident genetic divergence as revealed by branch length (Fig. 3), indicating that there might be six species within genus *Thermoleptolyngbya*. The 16S-23S ITS tree also detected clades corresponding to previously described genera supported by robust bootstrap values, namely *Alkalinema, Kovacikia, Leptolyngbya sensu stricto, Oculatella, Phormidesmis, Plectolyngbya*, and *Stenomitos*. Several strains corresponding to Cluster B and C in 16S rRNA tree (Fig. 2) were also included in the ITS tree and showed similar phylogenetic position or clustering as the 16S rRNA tree except for “*Leptolyngbya* sp.” PltLPT2.

**Fig. 3.**
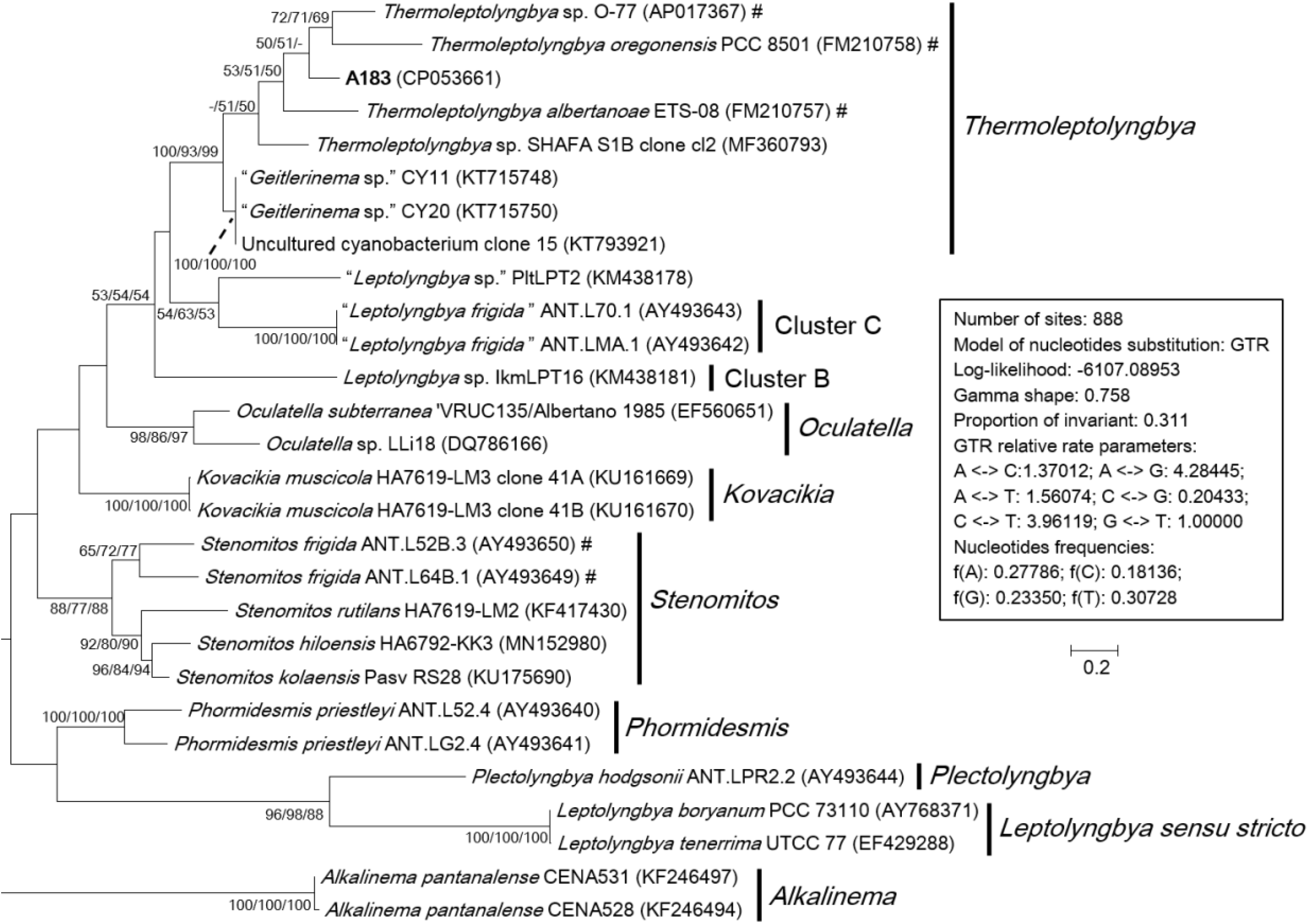
Maximum-likelihood phylogenetic tree of 16S-23S ITS sequences. The numbers at nodes refers to the support values of ML, MP, and NJ analysis, respectively. Only bootstrap values>50% (1000 nonparametric replications) are indicated at nodes. Genera or cluster are indicated by vertical bars. Scale bar=20% substitutions per site. Strains in “quotation marks” have uncertain genus name. Strains marked with # have been recently reclassified or proposed to be reclassified.

There are two primary reasons attributed to fewer sequences included in the 16S-23S ITS phylogenetic analysis than that in 16S rRNA analysis. Firstly, the 16S-23S ITS sequences are unavailable for many strains. In this study, only half of *Thermoleptolyngbya* strains had both of their 16S rRNA and 16S-23S ITS sequences determined, hindering further comprehensive taxonomic recognition. Secondly, more importantly, 16S-23S ITS marker is highly variable and is difficult to be aligned precisely if distantly related taxa were included in the database, affecting the outcome of phylogenetic reconstructions. It has been suggested that reliable phylogenies of 16S-23S ITS can be achieved by limiting the analysis to highly related taxa and with the support of secondary structure analysis (Johansen et al., 2011; Johansen et al., 2014). However, to ascertain the species identity more accurate analysis at molecular level is required, utilizing MLSA and eventually whole genome sequence comparison.

The 16S-23S ITS marker contains highly variable regions and highly conserved domains. The sequence identities are distinct when using full-length ITS or regions/domains individually, probably bringing about misleading information without secondary structure domain analysis. This speculation has been manifested by previous reports (Johansen et al., 2011; Sciuto & Moro, 2016). Additionally, the hyper-variable region of 16S-23S ITS marker is nearly neglected in the phylogenetic analysis since gapped positions are excluded. Those regions, however, may be informative at the species level. Therefore, secondary structure domain analysis of 16S-23S ITS is essential as a complement to ultimate taxonomy determination and is beneficial for better resolving the taxonomic status of problematic strains with inconsistent phylogenetic inferences between 16S rRNA and 16S-23S ITS.

### 3.4 Secondary structures of 16S-23S ITS

Hypothetical secondary structures of domains within ITS were estimated for strain A183 and representative strains from genus *Thermoleptolyngbya* in the 16S-23S ITS tree (Fig. 4). Excluding two highly conserved tRNAs from full-length ITS sequences, the length of the remaining ITS sequences varied greatly from 297 bp to 535 bp (Table 2). The remaining ITS sequence of A183 was the longest among *Thermoleptolyngbya* strains, 535 bp in length. Identical sequences were observed in conserved domains D3 (GGTTC), boxA (GAACCTTGAAAA), and D4 (CTGCATA) among all *Thermoleptolyngbya* strains. Conserved domain D2 showed four sequence types with slightly nucleotide difference, namely CTTCCAAACTAT in A183, O-77 and SHAFA S1B clone cl2, CTTCCAAGCTAG in ETS-08, CTTCCAAACTGT in CY11, and TTTCCAAACTAT in PCC 8501.

**Fig. 4.**
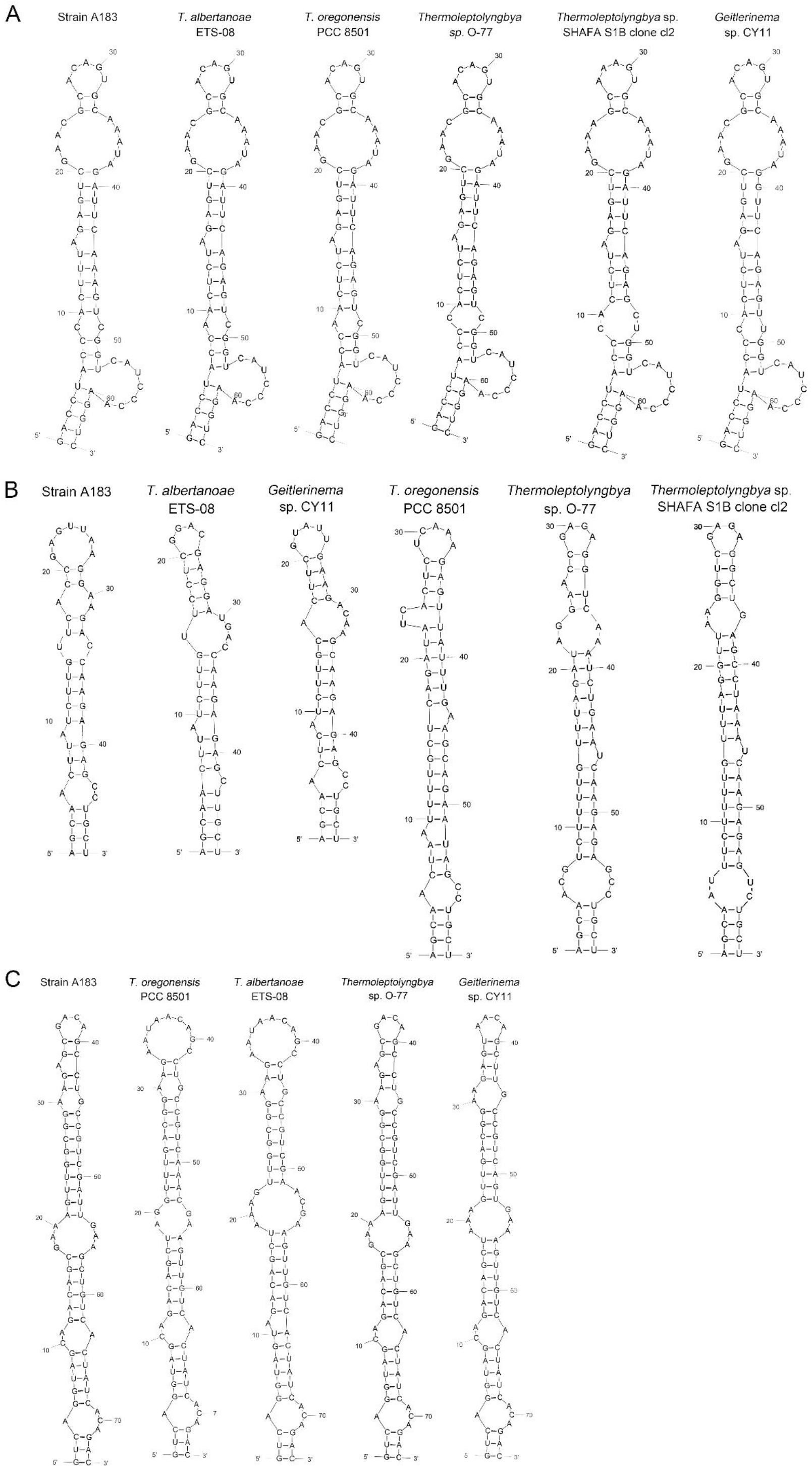
Predicted secondary structures of D1-D1’ helix, boxB and V3 helix of 16S-23S ITS of *Thermoleptolyngbya* strains.

**Table 2.** The length (bp) summary of regions within16S-23S ITS of strains studied. NA: not available

| Strain | ITS length<br>(tRNA removed) | D1-D1'<br>helix | D2 | D3 | boxA | D4 | V2<br>helix | boxB<br>helix | V3<br>helix |
| --- | --- | --- | --- | --- | --- | --- | --- | --- | --- |
| A183 | 535 | 64 | 12 | 5 | 12 | 7 | 218 | 48 | 74 |
| *Thermoleptolyngbya albertanoae* ETS-08 | 441 | 64 | 12 | 5 | 12 | 7 | 152 | 47 | 74 |
| *Thermoleptolyngbya oregonensis* PCC 8501 | 344 | 64 | 12 | 5 | 12 | 7 | 75 | 60 | 74 |
| *Thermoleptolyngbya* sp. O-77 | 486 | 64 | 12 | 5 | 12 | 7 | 211 | 60 | 74 |
| *Thermoleptolyngbya* sp. SHAFa S1B clone cl2 | 358 | 64 | 12 | 5 | 12 | 7 | 39 | 60 | NA |
| *Geitlerinema* sp. CY11 | 297 | 64 | 12 | 5 | 12 | 7 | 11 | 48 | 74 |

The inferred D1-D1’ helices were drawn in Fig. 4A. All helices exhibited the same length and nearly identical structures. However, nucleotide differences of D1-D1’ primary structures were found among strains. From basal stem (GACCU-AGGUC) onwards, all the helices were structured by 7 residue asymmetrical loop followed by 3 residue stem, 2 residue symmetrical loop followed by 5 residue stem, and one base left bulge followed by 5 residue stem, 9 residue asymmetrical loop, 2 residue stem and 5 residue asymmetrical loop. The only outlier of this structure is SHAFA S1B clone cl2 where 7 residue asymmetrical loop and 3 residue stem is followed by 4 residue stem and 4 residue symmetrical loop; all the remaining parts of the structure unchanged.

Hypothetical V2 helices were tremendously distinct among *Thermoleptolyngbya* strains (Fig. S2), and no common basal structure was found. The highly variable helices were attributed to the divergent primary structures of V2 regions. The longest V2 helix was found to be 218 residues in strain A183, followed by 211 residues in strain O-77, 152 residues in ETS-08, and 75 residues in PCC 8501, respectively (Table 2). The V2 helices of the remaining two strains were only 11 and 39 residues in length (Table 2).

The depicted boxB helices (Fig. 4B) indicated that a basal stem structure (AGCA-UGCU) was shared by all *Thermoleptolyngbya* strains. Although A183 had similar residue length to ETS-08 and CY11 (Table 2), the boxB helix structure of A183 was clearly distinct from that of the two strains. The boxB helix of A183 was mainly composed of a stem orderly fragmented by 3 residue asymmetrical loop, single base left bulge, 2 residue symmetrical loop, 3 residue asymmetrical loop, and terminated with 7 residue hairpin loop. The boxB helices of the other *Thermoleptolyngbya* strains (Fig. 4B) all terminated with hairpin loops variable in residues sequence and length, while the main stem structure of boxB helices considerably varied among strains in single base right bulge (in EST-08, PCC 8501, O-77, and SHAFA S1B clone cl2), single base left bulge (in EST-08, PCC 8501, O-77, and CY11), two base left bulge (in PCC 8501), asymmetrical loop (in EST-08, PCC 8501, SHAFA S1B clone cl2, and CY11), and symmetrical loop (in O-77).

The V3 helices shared a basal stem structure (GUC-GAC) among all *Thermoleptolyngbya* strains (Fig. 4C). The V3 helix of A183 comprised 2 asymmetrical loops, 2 symmetrical loops, single base left bulge, fragmented stems, and terminated with 4 residue hairpin loop. Although all the V3 helices showed the same helix length (Table 2), the structures (Fig. 4C) differed from each other in terms of bulge, loop and stem. Unfortunately, V3 helix of SHAFA S1B clone cl2 was not inferred due to incomplete sequences in this region.

In summary, the secondary structures of V2, boxB, and V3 undoubtedly differentiate A183 from the other *Thermoleptolyngbya* strains, whilst the structure of D1-D1’ domain remains conserved. The result of 16S-23S ITS secondary structure analysis is in agreement with the phylogenetic reconstructions of 16S-23S ITS, confirming the verification of A183 as a new species of *Thermoleptolyngbya*. The secondary structure analysis of V2, boxB, and V3 helices appear to be effective for species-level identification. Although the V2 helix was the most variable, it was the least taxonomic-informative in light of its high variability and absence in some cyanobacterial strains (Iteman et al., 2000; Sciuto & Moro, 2016). Although it was reported that D1-D1’ helix, compared to boxB and V3 helix, is more important for interspecies discrimination within a given genus (Perkerson et al., 2011; Vieira Vaz et al., 2015), it turned out to be not that effective in the case of *Thermoleptolyngbya*. The utilization of boxB and V3 for species distinction within *Thermoleptolyngbya* have been verified by the successful differentiation of *T. albertanoae* and *T. oregonensis* (Sciuto & Moro, 2016).

Interestingly, phylogenetic analysis and secondary structure analysis of 16S-23S ITS both indicated that *Thermoleptolyngbya* strains listed in Table 2 are probably different species to each other within the genus *Thermoleptolyngbya*, even though the phylogeny (Fig. 2) and sequence identity (Table S2) of 16S rRNA did not initially reveal such differentiation. Undoubtedly, detailed information regarding morphology and DNA sequence of more loci or complete genome would be essential for the taxonomic revision of SHAFA S1B clone cl2 and CY11within genus *Thermoleptolyngbya*.

### 3.5 Morphological investigation

The cell morphology of A183 indicated by light microscopy revealed straight, wavy, and occasionally bent trichomes (Fig. 5A) . The SEM and TEM (Fig. 5B - 5D) showed that trichomes of A183 were unbranched and composed of 80 - 120 elongated barrel-shaped cells, 1.30 – 1.60 μm in length and 1.05 - 1.10 μm in width. Constrictions were observed at the cross - walls of cells (Fig. 5B - 5D). Individual cells of the filaments were divided by centripetal invagination of the cell wall (Fig. 5E). Intracellular connections between vegetative cells were not observed (Fig. 5D). The TEM analysis also exhibited that the three to five thylakoids layers were located in parallel at the inner periphery of cells (Fig. 5C, 5D) and can be described as parietal according to recent classification (Mareš et al., 2019). Sheath, septum, phycobilisome, carboxysomes, cyanophycin granule, and polyphosphate bodies were present in the cytoplasm (Fig. 5C - 5F), and small lipid droplets were also observed (Fig. 5C).

**Fig. 5.**
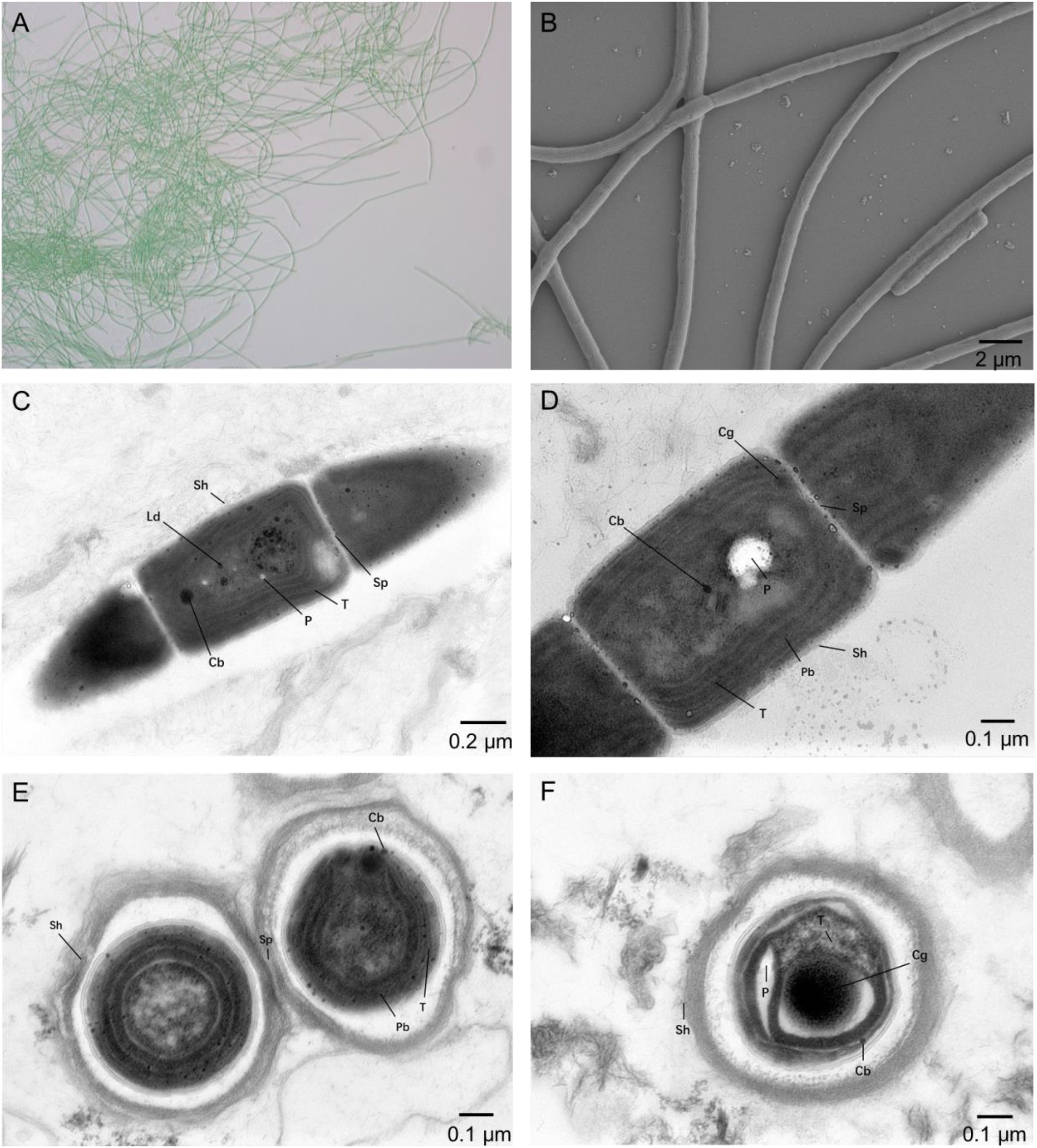
Micrographs of A183. **A**: light microscopy image. **B**: SEM image. **C - F**: TEM images. Abbreviations were as follows: Cb, carboxysome; Cg, cyanophycin granule; Cn, cyanophycin granule; Ld, lipid droplet; P, polyphosphate body; Pb, phycobilisome; Sh, sheath; Sp, septum; T, thylakoid membrane. Magnifications were 400× (A), 5,000x (B), 8,000 x (C) and 12,000 x (D-F).

The morphological characteristics of A183 showed certain similarity to that of other *Thermoleptolyngbya* strains (Sciuto & Moro, 2016). A183, together with ETS-08 and PCC 8501, were all blue-green filamentous cells and exhibited no vesicles. The sheath of A183 and PCC 8501 was unlayered, while ETS-08 showed multilayered sheath. There were granules observed at the cross walls of A183 and ETS-08 but no of PCC 8501. This observation reinforces the claim that on the morphology level alone it is impossible to make any final taxonomic conclusions.

### 3.6 Comparative genome analyses

To have an in-depth understanding of the A183 strain functioning in its ecosystem and to clarify the genetic features of the strain, the comparative genome analysis was performed between A183 and its relative *Thermoleptolyngbya* (O-77). Generally, the two strains shared a similar genome size, GC content, and the number of rRNA operons and tRNAs, but a big difference in CDS number (Table 1). Although fewer subsystems were observed in A183, subsystem category distribution was almost identical to both strains (Fig. S3). Analogously, GO analysis of all CDS showed a similar distribution of functional categories in both strains (Fig. S1).

An ortholog table (Table S3) was constructed based on all-against-all BLASTP alignment. As indicated in Fig. 1B, 4,344 genes of A183 were defined to be common to 4000 genes of O-77, accounting for 84% annotated genes of the A183 genome. The two strains shared 86.5% of their genomic segments, and these segments have 91.3% ANI value (Fig. 1C). This ANI value conformed to the suggested >96% intra-species and <83% inter-species ANI values (Jain & Rodriguez, 2018). Specific genes were also found in each strain, namely 822 genes in A183 and 1691 genes in O-77. Further, GO analysis of the specific genes showed that these genes were distributed in a wide range of functional category (Fig. S4). The result may suggest the specific ecological strategies of A183 relative to O-77.

No intact prophage region was detected in either strain. Three and eleven CRISPR arrays were identified in the genomes of A183 and O-77, respectively (Table S4). A higher number of spacers (597) was observed in the O-77 genome, approximately 3.5 times higher than that of A183. All direct repeats of the CRISPR arrays were 35 bp long in the A183 genome, while two kinds of length (34 and 35 bp long) in the O-77 genome. The A183 had only one CRISPR array with adjacent Cas genes. Although O-77 had two CRISPR arrays with Cas genes, one of them did not have a full set of genes encoding Cas system and was not assigned to a specific CRISPR-Cas system. The type III-D CRISPR-Cas system in both strains might confer resistance to foreign mobile genetic elements from bacteriophage or viruses (Makarova et al., 2011).

### 3.7 Thermotolerance

A183 was originally isolated from the hot spring with a temperature of 40.8°C (Tang et al., 2018a) and is capable of growing maximum at 50°C (Tang et al., 2018b). A survival strategy must be well-prepared for A183 to survive in these thermal environments. It is known that heat shock proteins (Hsps) are essential for maintaining and restoring protein homeostasis. In the A183 genome, dozens of genes were identified to encode heat shock proteins (Hsps), including the Hsp100, Hsp90, Hsp70 and Hsp60 family as well as small Hsps.

Hsps belonging to Hsp100 family may favor the refolding, disaggregation, removal of heat-damaged proteins. For instance, ATP-dependent proteases *ClpX*, associated with *ClpP*, promoted disassembly and degradation of heat-aggregated substrates (e.g. *FtsZ*) (Labreck et al., 2017). Homologues of *Clp* family were found in the A183 genome: *ClpB*, -*C*, -*P*, -*S*, -*X* (TS0414, TS0934, TS1039, TS1204-1206, TS1331, TS1444, TS1861-1862, TS2363, TS3986 and TS4547). An identical composition and distribution of *Clp* genes were found in O-77 genome (Table S1).

There was only one copy of *htpG* gene (TS0820) in A183, and so was O-77 (Table S1). The *htpG* protein of the Hsp90 family was suggested to be more of a general stress protein in that it played a role in several abiotic stresses (Hossain & Nakamoto, 2002; Hossain & Nakamoto, 2003). Particularly, the *htpG* primarily protected the photosynthetic machinery from heat stress in cyanobacteria (Takeshi et al., 2010).

Proteins of Hsp70 family appeared to be prosperous in A183 genome, mainly including *dnaK* (TS1830, TS2881, TS4129 and TS4674) and *dnaJ* (TS1379, TS2355, TS2878, TS4052, TS4637, TS4910 and TS5059). Multiple copies of *dnaK* and *dnaJ* genes have also been reported in other cyanobacteria (Rajaram et al., 2014). Nevertheless, it was suggested that *dnaK* and *dnaJ* proteins might function differently, and only some of them contributed to thermotolerance (Schneider, 2010; Duppre et al., 2011). Moreover, the gene *grpE* (TS2882) was found, the protein encoded by which might act as a cofactor of Hsp70 family and participate actively in response to heat shock by preventing the aggregation of stress-denatured proteins (Schneider, 2011). The homologues of Hsp70 family in O-77 showed a high similarity to that in A183 (Table S1).

Interestingly, two distinct *groEL* genes (TS0663, TS2946) of Hsp60 family, also referred to as the *groE* chaperone machinery, were identified in the A183 genome. One of them (TS2946, *groEL*-1) formed *groESL* operon together with the small Hsp *groES* (TS2947). The *Gloeobacter* PCC 7421 genome contains two *groEL* genes, both of which have *groES* immediately upstream of each *groEL*. Therefore, it was speculated that one of the two *groESL* operons had lost its *groES* during the evolutionary cyanobacterial diversification. Moreover, the relative expression of the two Hsp60 proteins against heat stress may be dependent on nitrogen status (Rajaram & Apte, 2008). The *HrcA* repressor system (TS4519) was also found in the A183 genome, which may negatively regulate the expression of *groE* genes (Nakamoto et al., 2003). A similar composition of *groE* genes was observed in the O-77 genome.

In addition to the genes mentioned above, genes encoding small Hsps and proteases were also identified in the A183 genome (Table S1) and might be involved in thermotolerance. For example, small *Hsp16* (TS0692), also referred to as *hspA*, may act as a chaperone and interact with dozens of proteins at high temperature, playing multiple roles ranging from protein folding to stabilization of thylakoid and periplasmic membranes (Basha et al., 2004); the *FtsH* protease (TS0606, TS1376, TS1741, TS1916 and TS2838) may be responsible for heat-induced degradation of photodamaged D1 protein by up-regulated expression of *FtsH* (Kamata et al., 2005). However, further investigations on the actual functions of these genes are necessary to elucidate the adaptation of A183 to high temperatures using other techniques such as RNA-Seq and targeted gene knock-outs.

### 3.8 Signal transduction

The two-component regulatory systems have been extensively found in cyanobacteria and elucidated for the perception of environmental stress and the subsequent transduction of stress signals (Los et al., 2010). In the A183 genome, 36 and 31 genes were identified to encode histidine kinases and response regulators, respectively (Table S1). The system composed of these genes is likely to play major roles in the core part of acclimation to changing environments. However, the scattered distribution of these genes in the A183 genome hindered the association of genes for histidine kinases with their respective cognate response regulators. This was in sharp contrast to cases in *E. coli* (Bourret & Silversmith, 2010) or *Bacillus subtilis* (Aguilar et al., 2001) that the genes for a single two-component system were organized into operons or located close one to another. Thus, investigation on sensors and their cognate regulators requires individual mutation on these genes. Similarly, the scattered genes encoding histidine kinases and response regulators were also found in the O-77 genome (Table S1).

The serine/threonine protein kinases (Spks) had similar purposes as two-component systems for signal transduction. A total of 17 genes were identified in A183 genome to encode Spks (Table S1). Unfortunately, the functions of only several Spks in *Synechocystis* have been characterized to date, such as *SpkA* and *SpkB* involved in the control of cell motility, and *SpkE* involved in the regulation of nitrogen metabolism (Zhang et al., 2006). Although the Spks proteins were conserved among the two *Thermoleptolyngbya* strains as revealed by their high similarity of amino acid sequences (87.61 - 96.71%, Table S3), the Spks proteins of *Thermoleptolyngbya* strains were quite divergent from these of other cyanobacterial strains. These putative Spks proteins in *Thermoleptolyngbya* are required to be more carefully investigated in future studies.

Different environmental conditions or developmental signals often cause major changes in transcription pattern by inducing a swap of sigma factors in the RNA polymerase holoenzyme (Los et al., 2010). The A183 genome inhabited seven genes encoding Group 2 sigma factor *sigD* (TS0188, TS0493, TS1286, TS1338, TS2057, TS3542 and TS4081) and two genes encoding Group 3 sigma factor *sigF* (TS1335 and TS3418). The *sigD* is the only *sig* gene that produced moderate amounts of transcripts in the dark and was not affected by any of stress treatments (Tuominen et al., 2003), suggesting its crucial role in transcription regulation particularly in adverse conditions. The exact function of *sigD* in cyanobacteria remains to be clarified.

In addition, seven genes encoding GGDEF/EAL domain proteins were observed in the A183 genome (Table S1). It was reported that GGDEF/EAL domain proteins function as diguanylate cyclase/phosphodiesterase that synthesizes/degrades cyclic di-GMP and participate in a cyclic-di-GMP signaling pathway that may regulate biofilm formation, motility, virulence, and cell cycle (Agostoni et al., 2013). A recent study showed that GGDEF/EAL domains were also involved in blue-light-induced cell aggregation in *Thermosynechococcus* BP-1 and NIES-2134 (Enomoto et al., 2015). Interestingly, *Thermosynechococcus* genomes have 9 - 13 genes encoding GGDEF/EAL domains, while hot-spring *Synechococcus* genomes (strain JA-3-3-Ab and JA-2-3Ba) only have four genes (Cheng et al., 2020). These data implied that the complexity of cyclic-di-GMP signaling pathways appeared to be distinct among thermotolerant strains.

### 3.9 Carbon assimilation

Cyanobacteria in hot springs have to deal with many environmental stresses, one of which is low CO_2_ solubility at high temperatures. The CO_2_-concentrating-mechanism (CCM) cyanobacteria have evolved can partially alleviate this problem by actively transporting and accumulating inorganic carbon (Ci: CO_2_ and HCO _3_^−^) for the sake of a satisfactory rate of CO_2_ fixation under carbon limiting concentrations (Price et al., 2008). The uptake of gaseous CO_2_ systems in cyanobacteria relied on NADPH dehydrogenase (NDH-1) complexes (Price et al., 2008). In the A183 genome, there were 19 NDH-1 genes encoding *ndhA, ndhB, ndhC, ndhE, ndhF, ndhG, ndhH, ndhI, ndhJ, ndhK, ndhL* and *ndhM*, respectively (Table S1). Except for *ndhE* and *ndhF*, the other NDH-1 genes were present in one copy. Meanwhile, several genes clustered together (*ndhE*-*ndhG*-*ndhI*-*ndhA*, TS1642-1645; *ndhJ*-*ndhK*-*ndhC*, TS2756-2758; *ndhF*-*ndhE*, TS4963-4964), while *ndhE* and *ndhF* composed of clusters coupled with low-affinity CO_2_ hydration proteins, namely *ndhF*-*ndhE*-*cphY* (TS4168-4170) and *ndhF*-*ndhE*-*cphX* (TS4275-4277). These gene clusters probably contribute to constitutively expressed NDH-1 complex involved in low-affinity CO_2_ uptake (Shibata et al., 2001). Similar clusters were also found in the O-77 genome.

Our previous study showed that A183 survived at the concentration exceeding 0.5M NaHCO_3_ (Tang et al., 2018b), indicating that A183 can assimilate bicarbonate and convert it to CO_2_ for photosynthesis. This was further evidenced by the HCO_3_^-^ uptake systems, as suggested by the genome analysis. First, a homolog (TS0333) of a low affinity, high flux, Na^+^-dependent bicarbonate transporter (*BicA*) was found in the A183 genome. Second, the A183 genome also harbored a homolog (TS2219) of *sbtA*, an inducible, high-affinity Na^+^-dependent bicarbonate transporter (Shibata et al., 2002). The different bicarbonate uptake systems might be flexibly utilized by A183 to meet the conditional demand for carbon gain. Interestingly, the *BCT1* bicarbonate transporter typical for many other cyanobacteria was not detected in the A183 genome, suggesting difference regarding a bicarbonate uptake mechanism from *Thermosynechococcus* BP-1 that lacks *sbtA* entirely but is equipped with *BCT1* (Price et al., 2008). In addition, three ABC-type bicarbonate transporters were observed in the A183 genome. The O-77 genome also possessed genes encoding *BicA* and *sbtA* and showed high protein similarities with A183 (94.99% and 95.56%, respectively), but the number of genes encoding ABC-type bicarbonate transporter was higher than that of A183 (9 versus 3).

### 3.10 Nitrogen assimilation

In the A183 genome, eight genes encoding nitrogenases were identified, including *nifB, nifR, nifH, nifO, nifW*, and *nifX* (Table S1). These genes encode proteins required for catalytic activity, Fe-Mo cofactor biosynthesis, and maturation and stability of the nitrogenase protein complex (Steunou et al., 2008). O-77 exhibited similar gene components regarding nitrogenases. These results suggested that both strains are nitrogen-fixing non-heterocystous cyanobacteria. N_2_ fixation is an energetically expensive metabolic reaction catalyzed by nitrogenase, which is inhibited by O_2_ generated during photosynthesis (Steunou et al., 2008). Therefore, A183 also exhibit alternative strategies in light of an economy of nitrogen assimilation.

Specific transporters are essential for organisms to concentrate ambient nitrogen sources within cells to survive in oligotrophic aquatic environments (Esteves-Ferreira et al., 2018). In the A183 genome, the ABC-type nitrate transport system (*nrtABCD*, TS4258-4261) was detected and formed an operon with two essential nitrogen-related genes located at both sides of *nrtABCD*, encoding Fd-nitrite reductase (*nirA*, TS4257) and Fd-nitrate reductase (*narB*, TS4262), respectively. This result was consistent with many freshwater cyanobacterial strains (Maeda et al., 2015). Furthermore, a complete gene set (*urtABCDE*) of ABC-type urea transport system and seven genes encoding urease (*ureA* – *ureG*) were observed (Table S1), suggesting the ability of A183 to import and utilize urea as a nitrogen source. In addition, two genes were identified as ammonium transporter (TS132, TS5152). The O-77 genome showed similar components of nitrogen-related transporters. The above results reinforced the importance of these transporters for cyanobacterial growth in oligotrophic environments and also implied that A183 can depend on multiple forms of nitrogen sources.

### 3.11 Sulfur assimilation

The isolation source of A183 was abundant in sulfur (Tang et al., 2018a). The analysis of the A183 genome reveals three transporter systems involved in sulfate uptake. First, the sulfate-thiosulfate permease (*sulT*), belonging to ABC-type transporter, comprised four subunits encoded by the *cysPTW* operon (TS1272-1274) and *cysA* (TS4980) that was located far away from the operon. The O-77 genome showed a different operon in terms of *cysPPTW* but a similar distribution of *cysA* gene. The organization of *cysPTW* operon in the *Thermoleptolyngbya* strains was different from that of *Thermosynechococcus* strains, such as BP-1 and PKUAC-SCTE542, resulting from the replacement of *cysP* by *sbpA* (Liang et al., 2019). The second sulfate permease in the A183 genome was *sulP* (TS1400), encoded by a single polypeptide and functioned as inorganic anion uptake carriers or anion:anion exchange transporters (Aguilar-Barajas et al., 2011). No homologue of *sulP* was identified in O-77 genome. Besides, sulfate can also be transported by the high-affinity ModABC molybdate transport system (Aguilar-Barajas et al., 2011), and the ModABC transporter in A183 genome was encoded by the *modABC* operon (TS2382-2384). Analogous ModABC transporter was also found in O-77 genome. These results indicated that there were different pathways of sulfur assimilation for A183. The predominant pathway and substrate specificity need to be experimentally clarified in future work.

## 4 Conclusions

The polyphasic approaches used in this study, including molecular, phylogenetic, ultrastructural, and morphological surveys, came up with a proposal of a new species, *Thermoleptolyngbya sichuanensis*. Although phylogenetic analysis and sequence identities of 16S rRNA showed a high similarity among *Thermoleptolyngbya* strains, phylogenetic reconstruction and secondary structures of 16S-23S ITS, together with the result of genome-scale average nucleotide identity (ANI), strongly indicate that A183 is a novel species within *Thermoleptolyngbya*. Comparative genome analysis revealed inconsistent genome structures of *Thermoleptolyngbya* strains. Moreover, genes related to thermotolerance, signal transduction, and carbon/nitrogen/sulfur assimilation were thoroughly analyzed for illustrating the ability of this strain to adapt to inhospitable niches in hot springs.

## Acknowledgements

This research was funded by National Natural Science Foundation of China (31970092, 32071480), Key Laboratory of Coarse Cereal Processing (Ministry of Agriculture and Rural Affairs, China) (2019CC12), and Tenure-Track Fund to MD.

## Supplementary Materials

**Table S1**. Annotations of protein-coding sequences in A183 and O-77 genome.

**Table S2**. Sequence identities of 16S rRNA gene between A183 and other *Thermoleptolyngbya* strains. Number in brackets indicated the pairwise alignment length. Strains are sorted by the order of identity from high to low. NA refers to data unavailable. Strains in “quotation marks” have uncertain genus name.

**Table S3**. Ortholog table. Constructed based on the all-against-all BLASTP alignments between A183 and O-77. The proteins are vertically shown in order of appearance in the genome of A183.

**Table S4**. CRISPR-Cas summary of *Thermoleptolyngbya* strains A183 and O-77.

**Fig. S1**. Gene ontology (GO) analysis and functional classification of genes in A183 and O-77 genomes identified in this study.

**Fig. S2**. Predicted secondary structures of V2 helix of 16S-23S ITS of *Thermoleptolyngbya* strains.

**Fig. S3**. Subsystem category distribution of A183 (A) and O-77 (B) based on the SEED databases.

**Fig. S4**. Gene ontology (GO) analysis and functional classification of specific genes in A183 and O-77 genomes identified in this study.

## References

Abed RM, Garcia-Pichel F, Hernández-Mariné M. 2002. Polyphasic characterization of benthic, moderately halophilic, moderately thermophilic cyanobacteria with very thin trichomes and the proposal of Halomicronema excentricum gen. nov., sp. nov. Archives of Microbiology 177: 361–370.

Agostoni M, Koestler BJ, Waters CM, Williams BL, Montgomery BL. 2013. Occurrence of cyclic di-GMP-Modulating output domains in cyanobacteria: an illuminating perspective. Mbio 4: 16–16.

Aguilar-Barajas E, Díaz-Pérez C, Ramírez-Díaz MI, Riveros-Rosas H, Cervantes C. 2011. Bacterial transport of sulfate, molybdate, and related oxyanions. BioMetals 24: 687–707.

Aguilar PS, Hernandez-Arriaga AM, Cybulski LE, Erazo AC, D. MD. 2001. Molecular basis of thermosensing: a two-component signal transduction thermometer in Bacillus subtilis. Embo Journal 20: 1681–1691.

Amarouche-Yala S, Benouadah A, Abd EOB, López-García P. 2014. Morphological and phylogenetic diversity of thermophilic cyanobacteria in Algerian hot springs. Extremophiles 18: 1035–1047.

Amin A, Ahmed I, Salam N, Kim B-Y, Singh D, Zhi X-Y, Xiao M, Li W-J. 2017. Diversity and distribution of thermophilic Bacteria in hot springs of Pakistan. Microbial Ecology 74: 116–127.

Andreote APD, Vaz M, Genuario DB, Barbiéro L, Fiore MF. 2014. Nonheterocytous cyanobacteria from Brazilian saline-alkaline lakes. Journal of Phycology 50: 675–684.

Arndt D, Grant JR, Marcu A, Sajed T, Pon A, Liang Y, Wishart DS. 2016. PHASTER: a better, faster version of the PHAST phage search tool. Nucleic Acids Research 44: W16–W21.

Basha E, Lee GJ, Breci LA, Hausrath AC, Buan NR, Giese KC, Vierling E. 2004. The identity of proteins associated with a small heat shock protein during heat stress in Vivo indicates that these chaperones protect a wide range of cellular functions. Journal of Biological Chemistry 279: 7566–7575.

Bourret RB, Silversmith RE. 2010. Two-component signal transduction. Current Opinion In Microbiology 13: 113–115.

Bravakos P, Kotoulas G, Skaraki K, Pantazidou A, Economou-Amilli A. 2016. A polyphasic taxonomic approach in isolated strains of Cyanobacteria from thermal springs of Greece. Molecular Phylogenetics and Evolution 98: 147–160.

Brito Â, Ramos V, Mota R, Lima S, Santos A, Vieira J, Vieira CP, Kaštovský J, Vasconcelos VM, Tamagnini P. 2017. Description of new genera and species of marine cyanobacteria from the Portuguese Atlantic coast. Molecular Phylogenetics and Evolution 111: 18–34.

Brown II, Bryant DA, Casamatta D, Thomas-Keprta KL, Sarkisova SA, Shen G, Graham JE, Boyd ES, Peters JW, Garrison DH. 2010. Polyphasic characterization of a thermotolerant siderophilic filamentous cyanobacterium that produces intracellular iron deposits. Applied and Environmental Microbiology 76: 6664–6672.

Bruno L, Dan I, Bellezza S, Albertano P. 2009. Cytomorphological and genetic characterization of troglobitic Leptolyngbya strains isolated from Roman Hypogea. Applied and Environmental Microbiology 75: 608–617.

Cheng Y-I, Chou L, Chiu Y-F, Hsueh H-T, Kuo C-H, Chu H-A. 2020. Comparative genomic analysis of a novel strain of Taiwan hot-spring cyanobacterium Thermosynechococcus sp. CL-1. Frontiers in Microbiology 11: 82.

Conesa A, Gotz S, Garcia-Gomez JM, Terol J, Talon M, Robles M. 2005. Blast2GO: a universal tool for annotation, visualization and analysis in functional genomics research. Bioinformatics 21: 3674–3676.

Debnath M, Singh T, Bhadury P. 2017. New records of Cyanobacterial morphotypes with Leptolyngbya indica sp. nov. from terrestrial biofilms of the Lower Gangetic Plain, India. Phytotaxa 316: 101–120.

Duppre E, Rupprecht E, Schneider D. 2011. Specific and promiscuous functions of multiple DnaJ proteins in Synechocystis sp. PCC 6803. Microbiology 157: 1269–1278.

Eckert EM, Fontaneto D, Coci M, Callieri C. 2014. Does a barcoding gap exist in prokaryotes? Evidences from species delimitation in cyanobacteria. Life 5: 50–64.

Enomoto G, Ni-Ni-Win, Narikawa R, Ikeuchi M. 2015. Three cyanobacteriochromes work together to form a light color-sensitive input system for c-di-GMP signaling of cell aggregation. Proceedings of the National Academy of Sciences of the United States of America 112: 8082–8087.

Esteves-Ferreira AA, Inaba M, Fort A, Araújo WL, Sulpice R. 2018. Nitrogen metabolism in cyanobacteria: metabolic and molecular control, growth consequences and biotechnological applications. Critical Reviews in Microbiology 44: 541–560.

Gaisin VA, Kalashnikov AM, Sukhacheva MV, Namsaraev ZB, Barhutova DD, Gorlenko VM, Kuznetsov BB. 2015. Filamentous anoxygenic phototrophic bacteria from cyanobacterial mats of Alla hot springs (Barguzin Valley, Russia). Extremophiles 19: 1067–1076.

Grissa I, Vergnaud G, Pourcel C. 2007. CRISPRFinder: a web tool to identify clustered regularly interspaced short palindromic repeats. Nucleic Acids Research 35: 52–57.

Guindon S, Dufayard J-F, Lefort V, Anisimova M, Hordijk W, Gascuel O. 2010. New algorithms and methods to estimate Maximum-Likelihood phylogenies: assessing the performance of PhyML 3.0. Systematic Biology 59: 307–321.

Heidari F, Zima J, Riahi H, Hauer T. 2018. New simple trichal cyanobacterial taxa isolated from radioactive thermal springs. Journal of the Czech Phycological Society 18: 137–149.

Hossain MM, Nakamoto H. 2002. HtpG plays a role in cold acclimation in cyanobacteria. Current Microbiology 44: 291–296.

Hossain MM, Nakamoto H. 2003. Role for the cyanobacterial htpG in protection from oxidative stress. Current Microbiology 46: 70–76.

Isabella Moro NR, Nicoletta La Rocca, Katia Sciuto, Patrizia Albertano, Laura Bruno, Andreoli C. 2010. Polyphasic characterization of a thermo-tolerant filamentous cyanobacterium isolated from the Euganean thermal muds (Padua, Italy). European Journal of Phycology 45: 143–154.

Iteman I, Rippka R, Tandeau DMN, Herdman M. 2000. Comparison of conserved structural and regulatory domains within divergent 16S rRNA-23S rRNA spacer sequences of cyanobacteria. Microbiology 146: 1275–1286.

Jain C, Rodriguez RL. 2018. High throughput ANI analysis of 90K prokaryotic genomes reveals clear species boundaries. Nature Communications 9: 5114.

Johansen JR, Kovacik L, Casamatta DA, Iková KF, Kaštovský J. 2011. Utility of 16S-23S ITS sequence and secondary structure for recognition of intrageneric and intergeneric limits within cyanobacterial taxa: Leptolyngbya corticola sp. nov. (Pseudanabaenaceae, Cyanobacteria). Nova Hedwigia 92: 283–302.

Johansen JR, Bohunická M, Lukešová A, Hrčková K, Vaccarino MA, Chesarino NM, Vis M. 2014. Morphological and molecular characterization within 26 strains of the genus Cylindrospermum (Nostocaceae, Cyanobacteria), with descriptions of three new species. Journal of Phycology 50: 187–202.

Kamata T, Hiramoto H, Morita N, Shen J-R, Mann NH, Yamamoto Y. 2005. Quality control of Photosystem II: an FtsH protease plays an essential role in the turnover of the reaction center D1 protein in Synechocystis PCC 6803 under heat stress as well as light stress conditions. Photochemical and Photobiological Sciences 4: 983–990.

Komárek J, Kaštovský J, Ventura S, Turicchia S, Šmarda J. 2009. The cyanobacterial genus Phormidesmis. Algological studies 129: 41–59.

Komarek J, Kaštovský J, Mares J, Johansen J. 2014. Taxonomic classification of cyanoprokaryotes (cyanobacterial genera), using a polyphasic approach. Preslia 86: 295–335.

Krzywinski MI, Schein JE, Birol I, Connors J, Gascoyne R, Horsman D, Jones SJ, Marra MA. 2009. Circos: An information aesthetic for comparative genomics. Genome Research 19: 1639–1645.

Kumar S, Stecher G, Tamura K. 2016. MEGA7: molecular evolutionary genetics analysis version 7.0 for bigger datasets. Molecular Biology and Evolution 33: 1870–1874.

Labreck CJ, Shannon M, Viola MG, Joseph C, Camberg JL. 2017. The protein chaperone ClpX targets native and non-native aggregated substrates for remodeling, disassembly, and degradation with ClpP. Frontiers in Molecular Bioences 4: 26.

Lacap DC, Barraquio W, Pointing SB. 2007. Thermophilic microbial mats in a tropical geothermal location display pronounced seasonal changes but appear resilient to stochastic disturbance. Environmental Microbiology 9: 3065–3076.

Li H, Durbin R. 2009. Fast and Accurate Short Read Alignment with Burrows-Wheeler Transform. Bioinformatics 25: 1754–1760.

Liang Y, Tang J, Luo Y, Kaczmarek MB, Li X, Daroch M. 2019. Thermosynechococcus as a thermophilic photosynthetic microbial cell factory for CO_2_ utilisation. Bioresource Technology 278: 255–265.

Los DA, Zorina A, Sinetova M, Kryazhov S, Mironov K, Zinchenko VV. 2010. Stress sensors and signal transducers in cyanobacteria. Sensors 10: 2386–2415.

Lowe TM, Eddy SR. 1997. tRNAscan-SE: a program for improved detection of transfer RNA genes in genomic sequence. Nucleic Acids Research 25: 955–964.

Mackenzie R, Pedrós-Alió C, Díez B. 2012. Bacterial composition of microbial mats in hot springs in Northern Patagonia: Variations with seasons and temperature. Extremophiles 17: 123–136.

Maeda S-i, Murakami A, Ito H, Tanaka A, Omata T. 2015. Functional characterization of the FNT family nitrite transporter of marine Picocyanobacteria. Life 5: 432–446.

Makarova KS, Haft DH, Barrangou R, Brouns SJ, Charpentier E, Horvath P, Moineau S, Mojica FJ, Wolf YI, Yakunin AF, van der Oost J, Koonin EV. 2011. Evolution and classification of the CRISPR-Cas systems. Nature reviews Microbiology 9: 467–477.

Mareš J, Strunecky O, Bučinská L, Bučinská B, Wiedermannová J. 2019. Evolutionary patterns of thylakoid architecture in cyanobacteria. Frontiers in Microbiology 10: 277.

Mikheeva LE, Karbysheva EA, Shestakov SV. 2013. The role of mobile genetic elements in the evolution of cyanobacteria. Russian Journal of Genetics 3: 91–101.

Miscoe LH, Johansen JR, Kociolek JP, Lowe RL, Vaccarino MA, Pietrasiak N, Sherwood AR. 2016. The diatom flora and cyanobacteria from caves on Kauai, Hawaii. II. Novel cyanobacteria from caves on Kauai, Hawaii. Bibliotheca Phycologica: 152.

Nakamori H, Yatabe T, Yoon KS, Ogo S. 2014. Purification and characterization of an oxygen-evolving photosystem II from Leptolyngbya sp. strain O-77(enzymology, protein engineering, and enzyme technology). Journal of Bioscience and Bioengineering 118: 119–124.

Nakamoto H, Suzuki M, Kojima K. 2003. Targeted inactivation of the hrcA repressor gene in cyanobacteria. FEBS Letters 549: 57–62.

Niclas E, Cameron CR, Williamh G. 2010. 16S rRNA gene heterogeneity in the filamentous marine cyanobacterial genus Lyngbya. Journal of Phycology 46: 591–601.

Oren A, Ionescu D, Hindiyeh M, Malkawi H. 2008. Morphological, phylogenetic and physiological diversity of cyanobacteria in the hot springs of Zerka Ma. BioRisk 3: 20–23.

Pawan, K., Dadheech, Huda, Mahmoud, Kiplagat, KotutLothar, Krienitz. 2012. Haloleptolyngbya alcalis gen. et sp. nov., a new filamentous cyanobacterium from the soda lake Nakuru, Kenya. Hydrobiologia 691: 269–283.

Peng X, Xu H, Jones B, Chen S, Zhou H. 2013. Silicified virus-like nanoparticles in an extreme thermal environment: implications for the preservation of viruses in the geological record. Geobiology 11: 511–526.

Perkerson RB, Johansen JR, Kovácik L, Brand J, Kaštovský J, Casamatta DA. 2011. A unique Pseudanabaenalean (cyanobacteria) genus Nodosilinea gen. nov. based on morphological and molecular data. Journal of Phycology 47: 1397–1412.

Price GD, Badger MR, Woodger FJ, Long BM. 2008. Advances in understanding the cyanobacterial CO_2_-concentrating-mechanism (CCM): functional components, Ci transporters, diversity, genetic regulation and prospects for engineering into plants. Journal of Experimental Botany 59: 1441–1461.

Pruitt KD, Tatusova T, Klimke W, Maglott DR. 2009. NCBI Reference Sequences: current status, policy and new initiatives. Nucleic Acids Research 37: D32–D36.

Rajaram H, Apte SK. 2008. Nitrogen status and heat-stress-dependent differential expression of the cpn60 chaperonin gene influences thermotolerance in the cyanobacterium Anabaena. Microbiology 154: 317–325.

Rajaram H, Chaurasia AK, Apte SK. 2014. Cyanobacterial heat-shock response: role and regulation of molecular chaperones. Microbiology 160: 647–658.

Rodriguez-R LM, Santosh G, Harvey WT, Ramon R-M, Tiedje JM, Cole JR, Konstantinidis KT. 2018. The Microbial Genomes Atlas (MiGA) webserver: taxonomic and gene diversity analysis of Archaea and Bacteria at the whole genome level. Nuclc Acids Research: W282-W288.

Roy, Mackenzie, Carlos, Pedrós-Alió, Beatriz, Díez. 2013. Bacterial composition of microbial mats in hot springs in Northern Patagonia: variations with seasons and temperature. Extremophiles 17: 123–136.

Schneider D. 2010. Similarities and singularities of three DnaK proteins from the cyanobacterium Synechocystis sp. PCC 6803. Plant And Cell Physiology 51: 1210–1218.

Schneider D. 2011. Thermostability of two cyanobacterial GrpE thermosensors. Plant And Cell Physiology 52: 1776–1785.

Sciuto K, Moro I. 2016. Detection of the new cosmopolitan genus Thermoleptolyngbya (Cyanobacteria, Leptolyngbyaceae) using the 16S rRNA gene and 16S–23S ITS region. Molecular Phylogenetics and Evolution 105: 15–35.

Shalygin S, Shalygina R, Redkina V, Gargas C, Johansen J. 2020. Description of Stenomitos kolaenensis and S. hiloensis sp. nov. (Leptolyngbyaceae, Cyanobacteria) with an emendation of the genus. Phytotaxa 440: 108–128.

Shibata M, Ohkawa H, Kaneko T, Fukuzawa H, Tabata S, Kaplan A, Ogawa T. 2001. Distinct constitutive and low-CO_2_ induced CO_2_ uptake systems in cyanobacteria: Genes involved and their phylogenetic relationship with homologous genes in other organisms. Proceedings of the National Academy of Sciences 98: 11789–11794.

Shibata M, Katoh H, Sonoda M, Ohkawa H, Shimoyama M, Fukuzawa H, Kaplan A, Ogawa T. 2002. Genes essential to sodium-dependent bicarbonate transport in cyanobacteria: function and phylogenetic analysis. Journal of Biological Chemistry 277: 18658–18664.

Steunou A-S, Jensen SI, Brecht E, Becraft ED, Bateson MM, Kilian O, Bhaya D, Ward DM, Peters JW, Grossman AR, Kühl M. 2008. Regulation of nif gene expression and the energetics of N_2_ fixation over the diel cycle in a hot spring microbial mat. The ISME journal 2: 364–378.

Strunecký O, Kopejtka K, Goecke F, Tomasch J, Lukavský J, Neori A, Kahl S, Pieper DH, Pilarski P, Kaftan D, Koblížek M. 2019. High diversity of thermophilic cyanobacteria in Rupite hot spring identified by microscopy, cultivation, single-cell PCR and amplicon sequencing. Extremophiles 23: 35–48.

Takeshi, Sato, Shun, Minagawa, Erika, Kojima, Naoki, Okamoto, Hitoshi, Nakamoto. 2010. HtpG, the prokaryotic homologue of Hsp90, stabilizes a phycobilisome protein in the cyanobacterium Synechococcus elongatus PCC 7942. Molecular Microbiology 76: 576–589.

Tang J, Du L-M, Liang Y-M, Daroch M. 2019. Complete genome sequence and comparative analysis of Synechococcus sp. CS-601 (SynAce01), a cold-adapted cyanobacterium from an oligotrophic Antarctic habitat. International Journal of Molecular Sciences 20: 152.

Tang J, Liang Y, Jiang D, Li L, Luo Y, Shah MMR, Daroch M. 2018a. Temperature-controlled thermophilic bacterial communities in hot springs of western Sichuan, China. BMC Microbiology 18: 134.

Tang J, Jiang D, Luo Y, Liang Y, Li L, Shah MMR, Daroch M. 2018b. Potential new genera of cyanobacterial strains isolated from thermal springs of western Sichuan, China. Algal Research 31: 14–20.

Taton A, Wilmotte A, Smarda J, Elster J, Komarek J. 2011. Plectolyngbya hodgsonii: a novel filamentous cyanobacterium from Antarctic lakes. Polar Biology 34: 181–191.

Taton A, Grubisic S, Ertz D, Hodgson DA, Piccardi R, Biondi N, Tredici MR, Mainini M, Losi D, Marinelli F. 2010. Polyphasic study of Antarctic cyanobacterial strains. Journal of Phycology 42: 1257–1270.

Tuominen I, Tyystjärvi E, Tyystjärvi T. 2003. Expression of primary sigma factor (PSF) and PSF-Like sigma factors in the cyanobacterium Synechocystis sp. strain PCC 6803. Journal of Bacteriology 185: 1116–1119.

Varani AM, Siguier P, Gourbeyre E, Charneau V, Chandler M. 2011. ISsaga is an ensemble of web-based methods for high throughput identification and semi-automatic annotation of insertion sequences in prokaryotic genomes. Genome Biology 12: 1–9.

Vaz MGMV, Genuário DB, Andreote APD, Malone CFS, Sant’Anna CL, Barbiero L, Fiore MF. 2015. Pantanalinema gen. nov. and Alkalinema gen. nov.: novel pseudanabaenacean genera (Cyanobacteria) isolated from saline–alkaline lakes. International Journal of Systematic and Evolutionary Microbiology 65: 298–308.

Vieira Vaz MG, Genuário DB, Andreote AP, Malone CF, Sant’Anna CL, Barbiero L, Fiore MF. 2015. Pantanalinema gen. nov. and Alkalinema gen. nov.: novel pseudanabaenacean genera (Cyanobacteria) isolated from saline-alkaline lakes. International Journal of Systematic and Evolutionary Microbiology 65: 298.

Vincent L, Jean-Emmanuel L, Olivier G. 2017. SMS: Smart Model Selection in PhyML. Molecular Biology and Evolution 34: 2422–2424.

Walker BJ, Abeel T, Shea T, Priest M, Abouelliel A, Sakthikumar S, Cuomo CA, Zeng Q, Wortman J, Young SK. 2014. Pilon: an integrated tool for comprehensive microbial variant detection and genome assembly improvement. Plos One 9: e112963.

Wang Z, Xiao P, Song G, Li Y, Li R. 2015. Isolation and characterization of a new reported cyanobacterium Leptolyngbya bijugata coproducing odorous geosmin and 2-methylisoborneol. Environmental Science And Pollution Research 22: 12133–12140.

Ye J, Fang L, Zheng H, Zhang Y, Chen J, Zhang Z, Wang J, Li S, Li R, Bolund L, Wang J. 2006. WEGO: a web tool for plotting GO annotations. Nucleic Acids Research 34: 293–297.

Zammit G, Billi D, Albertano P. 2012. The subaerophytic cyanobacterium Oculatella subterranea (Oscillatoriales, Cyanophyceae) gen. et sp. nov.: a cytomorphological and molecular description. European Journal of Phycology 47: 341–354.

Zhang CC, Jang J, Sakr S, Wang L. 2006. Protein phosphorylation on Ser, Thr and Tyr residues in cyanobacteria. Journal of Molecular Microbiology and Biotechnology 9: 154–166.

Zimin AV, Marçais G, Puiu D, Roberts M, Salzberg SL, Yorke JA. 2013. The MaSuRCA genome assembler. Bioinformatics 29: 2669–2677.

Zuker M. 2003. Mfold web server for nucleic acid folding and hybridization prediction. Nucleic Acids Research 31: 3406–3415.

